# Ecology of Chagas disease vector *Triatoma sanguisuga* in Delaware: distribution, habitat, host use and *Trypanosoma cruzi* infection

**DOI:** 10.64898/2026.11.12.766167

**Authors:** Alexander R. Kelley, Hunt Kinnaird, Hanna Cortes, Grace Ferry, Alexandra Hemelt, Jillian Tunstall, Jennifer K. Peterson

## Abstract

Chagas disease is a vector-borne disease caused by the parasite *Trypanosoma cruzi* and transmitted by triatomine bugs. The disease is increasingly recognized as an emerging public health threat in the United States (US). Little is known about the Chagas disease system in the Mid-Atlantic US despite the presence of a native *T. cruzi* vector species, *Triatoma sanguisuga*. The species is widespread throughout the eastern US, and the Mid-Atlantic comprises the northernmost portion of its geographic range. *Triatoma sanguisuga* has been implicated in autochthonous *T. cruzi* transmission to humans in other parts of the US but has never been formally studied in the Mid-Atlantic. To fill this knowledge gap and increase public health preparedness, we investigated *T. sanguisuga* in Delaware and eastern Maryland. We aimed to collect baseline information about the species, including its habitat, feeding patterns, and *T. cruzi* infection rates across land use intensities.

We collected 151 *T. sanguisuga* specimens in all three Delaware counties and two counties in Maryland, and identified standing dead trees as an important local habitat for the species. Twenty specimens (13.2%) were submitted by community members who had found the bugs in or around human homes and businesses. We detected *T. cruzi* infection in 23 of 150 (15.3%) specimens and identified 24 vertebrate host species, including humans, from 95 specimens. We detected a stage-structured blood feeding pattern wherein juvenile bugs (‘nymphs’) were reptile-associated (which are refractory *T. cruzi* hosts) and adult bugs were mammal-associated (which are competent *T. cruzi* hosts), which was reflected in a significant difference in *T. cruzi* infection prevalence between adults and nymphs. Our findings reveal the multiscale network that sustains *T. cruzi* transmission in Delaware while also providing practical public health information on a local vector-borne disease system.

## Introduction

Chagas disease is a vector-borne zoonotic disease caused by infection with the protozoan parasite of mammals, *Trypanosoma cruzi.* When left untreated in humans, *T. cruzi* infection is chronic and can lead to serious cardiac or gastrointestinal alterations [1]. The World Health Organization (WHO) estimates that eight million people are infected with *T. cruzi* worldwide and considers more than 100 million to be at risk of infection. There are an estimated 10,000 deaths each year as a result of Chagas disease [2].

*Trypanosoma cruzi* is transmitted between mammal hosts by triatomine bugs (Hemiptera: Reduviidae). There are 159 extant triatomine bug species, most of which are concentrated in the New World [3–6]. Eleven triatomine bug species have been described in the United States (US), where they are reported as ranging up to approximately the 42^nd^ parallel north [7]. The widest-ranging species in the US is *Triatoma sanguisuga*; it has been reported in 26 states from the Rocky Mountains to the eastern seaboard as far north as New Jersey [8–12].

*Triatoma sanguisuga* is a competent vector of *T. cruzi*, and has been found infected with the parasite with prevalences ranging from 20% to 60% [13–15]. The insect has been documented infected with *T. cruzi* inside homes [13,16,17] and implicated in confirmed cases of autochthonous Chagas disease in Tennessee, Louisiana, and Mississippi [18–20]. Reports of domestic infestations of *T. sanguisuga* are rare, but adults do sporadically fly into human homes and feed on resident humans and animals. Indeed, the species was first described in Georgia by LeConte in 1855 as, ‘sucking the blood of mammals, particularly of children’ [21–23]. More recent blood meal studies confirm that these behaviors still occur; *T. sanguisuga* blood meals have been detected from several domestic and agricultural animal species as well as from humans [13,16,24,25].

Despite its broad distribution and epidemiological relevance, *T. sanguisuga* remains poorly studied outside the southern US, particularly at northern latitudes with cooler climates. The species is found as far north as Illinois in the Midwest and New Jersey on the east coast; these regions have not only cooler climates than the southern US but also key differences in wildlife communities, topography, and vegetation [26], which could influence epidemiologically relevant features of *T. sanguisuga* ecology such as blood feeding, host seeking, and parasite competence.

Here, we collected baseline eco-epidemiological information on the occurrence, *T. cruzi* infection prevalence, host feeding, and habitat use of *T. sanguisuga* in the Mid-Atlantic US, beginning with the state of Delaware and adjacent portions of Maryland. The Mid-Atlantic region (comprising Delaware, Maryland, New Jersey, Pennsylvania, Virginia, West Virginia and Washington DC [27]) constitutes the extreme northeastern edge of *T. sanguisuga*’s geographic distribution and by extension that of the subfamily Triatominae. *Triatoma sanguisuga* has been documented in the Mid-Atlantic for over a century, but has not been the subject of focused scientific investigation in the region [17,28–35]. Our aim was to provide fundamental epidemiological information on *T. sanguisuga* in temperate environments as a first step toward enabling Mid-Atlantic citizens to make evidence-based decisions when encountering this important disease vector.

## Methods

### Overview

To investigate the geographic distribution, *Trypanosoma cruzi* infection, habitat use, and blood-feeding behavior of *Triatoma sanguisuga* in Delaware (DE), we sampled for *T. sanguisuga* across different land use types. Each *T. sanguisuga* specimen underwent *T. cruzi* infection detection and blood meal analysis. Triatomine sampling took place between May and September of 2024 (‘Year 1’) and June through August of 2025 (‘Year 2’), which has been found to be the active period for *T. sanguisuga* in other regions [36,37]. Given the dearth of local *T. sanguisuga* data, our Year 1 sampling design was based on prior studies of other triatomine species in cooler climates [38,39], where triatomine distribution is clustered and heterogeneous. We used historical *T. sanguisuga* occurrence records to identify potential sites and sampled them repeatedly used multiple collection methods in conjunction with community outreach to detect the species. In Year 2, we used data collected in Year 1 to refine our methods and carry out a more intensive, targeted approach, as described below.

### Permitting

Sampling was carried out under the Delaware Department of Natural Resources and Environmental Control (DNREC) DE Scientific Collecting Permit Nos. 2025-WSC-031 and 2026-WSC-035. Residential and privately owned sites, state parks, city parks and state forests were sampled with written permission from the owner or corresponding property manager. Maryland specimens were submitted to us unsolicited by residents who found them in their homes or businesses.

### Study sites

We actively sampled 28 sites located throughout the three DE counties (eight in Year 1 and 20 in Year 2), which varied in size from private residential properties (∼1/2 acre) to state parks or wildlife refuges (100s of acres). We aimed to sample each site a minimum of four times Year 1 sites were selected via the following selection criteria: (i) at least one *T. sanguisuga* specimen occurrence (determined by reports on inaturalist.com and bugguide.org and specimen metadata from the University of Delaware entomological collection); (ii) sampling was feasible, both in terms of accessibility and permission. If these criteria were met, we then considered the land use intensity (LUI) of the site with the aim of sampling across different epidemiological contexts [40]. We classified sites as high, medium, or low LUI. High LUI sites had frequent human presence and significant landscape alteration, such as residential properties, commercial buildings, and small urban parks. Medium LUI sites had intermittent human presence and a mixed landscape of natural and artificial habitat, such as outdoor recreation areas with trails and campsites. Low LUI sites had a largely unaltered landscape of natural habitat where human presence was rare, such as wildlife refuges or nature preserves. Community submission locations were also assigned an LUI designation.

In Year 2, we used data collected in Year 1 to refine our study site selection criteria to (i) regions of DE that we had not yet sampled; (ii) sampling was feasible, both in terms of accessibility and permission; and (iii) sampling evenly across the three LUIs. We discarded the criterion of previous *T. sanguisuga* sighting.

### Collection methods

We collected *T. sanguisuga* specimens using four methods: manual searching of standing dead trees (snags), light sheet traps, bucket traps (Year 1 only), and community submissions, as follows:

- **Manual searching**: pieces of loose bark were peeled off standing dead trees and the underside of the bark, outer surface of exposed trunk, and tree cavities were examined for triatomine bugs. The circumference of a snag was debarked in this manner, limited in height to the unaided reach of team members, as snags could not be safely climbed.
- **Light sheet traps**: a twin sized (66in by 96in) white bed sheet was hung vertically from a freestanding metal frame of the same size and staked to the ground. The sheet was illuminated by two lights: an ADJ Black-24BLB 20W fluorescent tube style light that emitted UV light at a 365nm wavelength and an Entoquip Entolight 32W LED light (www.entoquip.com) that emitted UV light at both 365nm and 395nm. The ADJ light was hung from the top bar of the metal frame, contacting the top of the sheet while the Entolight light was hung from a freestanding shepherd’s hook embedded in the ground approximately two feet in front of the sheet. Both lights were powered by a single Progeny 300W Portable Power Station.
- **Ultraviolet light bucket traps:** automated light traps that used artificial light to intercept nocturnal flying insects in a manner similar to that used in light sheet traps [41]. Rather than landing on an illuminated surface, insects approaching the light collided with a clear plastic barrier and dropped through a funnel into a collection chamber [42]. These traps successfully collected other *Triatoma* species in Texas [43]. Bucket traps consisted of two nine-light UV LED strips mounted on two clear plastic vanes, wired to a top unit holding 9V batteries. This apparatus rested on a funnel over the open end of a 10-gallon bucket. Bucket traps were placed at study sites for 24 or 48 hours after which they were recollected and their contents were inspected for triatomines. These traps were used only in Year 1.
- **Community submissions**: community members in Delaware and eastern Maryland submitted *Triatoma sanguisuga* specimens that they encountered unexpectedly in their homes, offices, or businesses. These individuals located us through word of mouth, the internet, and an educational flyer distributed at study sites and community outreach events. The flyer contained information about Chagas disease, pictures to aid in identifying *T. sanguisuga*, and contact information for specimen submission. At these events we informed community members about Chagas disease and encouraged the safe collection and disposal of triatomine specimens. We also provided an email address to contact with questions.

### Field sampling protocol

As mentioned, sample sites varied in size from private residential properties (∼1/2 acre) to state parks or wildlife refuges (100s of acres). At sites for which two light sheets illuminated the entire site and manual searching of the whole site was feasible, we sampled at the same location within the site four times. For sites comprising several hundred or thousand acres, we attempted to sample evenly across the site throughout the four site visits. When light sheet collections were successful in larger sites, we sampled the same location repeatedly, given the low density and clustered nature of *T. sanguisuga* distribution. When light sheets were unsuccessful, we chose a new location upon the next visit according to the criteria listed above.

To maximize light sheet performance, we chose dark areas away from competing light sources with minimal dense vegetation or other obstacles that prevent the light from illuminating a wider area. At high LUI sites we sampled as close as possible to residences or commercial buildings to collect triatomines that may have fed on humans or their animals. Similarly, in medium LUI sites we sampled near recreational infrastructure such as cabins or trails. Conversely, in low LUI sites we aimed to sample as far from human influence as possible to capture sylvatic context.

The sampling protocol consisted of 60 minutes of manual searching in which team members searched all snags within a 100 meter radius of the sampling location. After completing manual searching, we deployed light sheets. We utilized one or two light sheets depending on the size of the site and the number of team members available to monitor the sheets. When using two light sheets, we separated them such that light from one light sheet was not visible from the other sheet. We turned lights on at sundown, defined as the beginning of civil twilight based on a timetable from the US Naval Observatory [44]. Lights remained on for at least two hours [45–48], during which each light sheet was observed continuously by one or two team members.

### Metadata collection

At each sampling location, we recorded the date, arrival time, and GPS coordinates of the site. Immediately before and after the sampling period, we recorded temperature, humidity, and wind speed using a LaCrosse Technology Wireless Temperature and Humidity Station and a BTMeter BT-100 Pro-Anemometer. We did not sample in the rain or during windstorms or electrical storms. If a wind or a rainstorm began during light sheet sampling, we would end early or truncate the protocol and perform only manual searching with no light sheets. For each specimen collected (field collected and community submissions), we recorded the date and time, geographic coordinates, and method of collection.

### Specimen processing and microscopy

For each specimen, we recorded its stage (nymph or adult), sex (adults only), and weight. If specimens were alive, we extracted excreta to check for *T. cruzi* forms microscopically. In this procedure, we expressed excrement from live bugs by applying light pressure using forceps to the posterior end of the abdomen at the approximate location of the rectum until fluid was expressed from the anus onto a microscope slide. The fluid was then mixed on the slide with approximately 5μl of phosphate buffer saline and examined under a compound microscope at 40X magnification. We identified *T. cruzi* forms by morphology and flagellum movement. Dead specimens did not undergo this examination. In some cases, we were unable to express fluid from live specimens that were in a starved state. After this procedure, insects were stored at - 20°C until further analysis.

### DNA extraction

DNA was extracted from whole bugs using phenol-chloroform and Qiagen blood and tissue kits. Bugs were immersed in a solution of 5% sodium hypochlorite before extraction to eliminate DNA on the surface of the bug. DNA from specimens collected in Year 1 was extracted using a standard Phenol-Chloroform (PCI) extraction technique [49] due to its tendency to yield higher DNA concentrations when compared to commercial kits, which was our priority when extracting insect, parasite, and blood meal DNA from a single specimen [50]. However, the resulting DNA required additional cleaning to reduce RNA and pigments prior to Illumina sequencing, so Year 2 DNA was extracted using Qiagen DNeasy Blood & Tissue Kits following manufacturer’s instructions. DNA was extracted from each specimen in groups of no more than 10 extractions at once. Each group of 10 extractions also included a negative control with reagents only. After DNA extraction, DNA concentration was measured in each sample using a Qubit 4 Fluorometer (Invitrogen). Samples were then diluted to a standard concentration of 5ng/μL and stored at -20°C.

### RT-PCR detection of *Trypanosoma cruzi*

We tested each specimen for *T. cruzi* infection using an RT-PCR assay as described in Moreira et al. [51]. Briefly, the assay consisted of a multiplex TaqMan real time PCR using primers targeting a repeating sequence of nuclear satellite DNA (SAT-DNA) in the *T. cruzi* genome (Table 1). Primers targeting a sequence of the triatomine 12S rRNA gene were used as an internal amplification control (IAC). The reaction was carried out using 2X TaqMan Fast Advanced Master Mix (Applied Biosystems) in a final volume of 20μL containing 1μL of extracted DNA solution at a concentration of 5ng/μL. The cycling conditions were as follows: 50°C for 2 minutes, 95°C for 10 minutes, then 40 cycles of 95°C for 15 seconds and 58°C for 1 minute. The reaction was run in a QuantStudio3 Real-Time PCR System (Applied Biosystems).

**Table 1.**
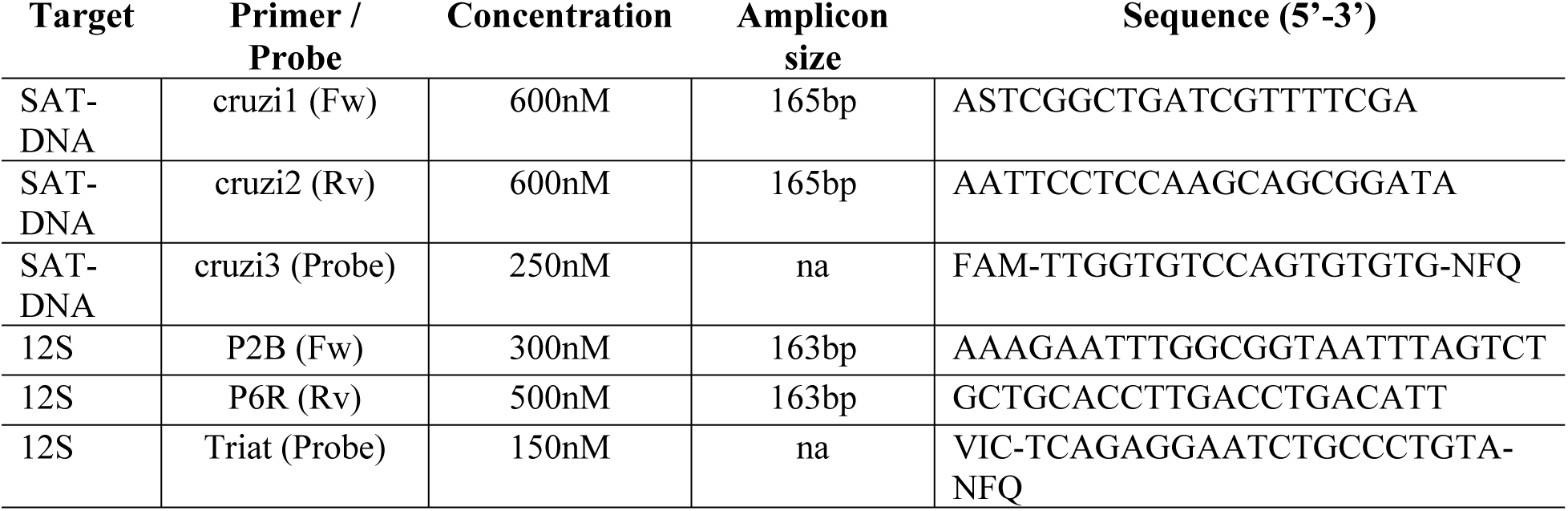
Primers and probes used to detect *T. cruzi* infection in *T. sanguisuga* specimens. Final reaction concentrations, amplicon lengths, and sequences for primers and probes in the diagnostic RT-PCR assay. In sequence notation, FAM and VIC denote fluorescent dyes, NFQ denotes a non-fluorescent quencher molecule, S denotes a degenerate G/C base following IUPAC nomenclature[52]

DNA from *T. cruzi*-positive specimens was run in a follow up RT-PCR assay to determine the *T. cruzi* discrete typing unit (DTU), as described in Cura et al. [53]. This assay differentiated between six *T. cruzi* DTUs (TcI - TcVI) but not TcBat [53]. Primer and sequence details for this assay are found in the supplementary materials (Table S1).

### Blood meal analysis

To identify vertebrate host species on which each *T. sanguisuga* had fed, we employed next generation sequencing using a metabarcoding approach following Kieran *et al.* [54]. This assay amplified a universally conserved 145bp fragment of the vertebrate 12S rRNA gene using the primers 12S-6F (5′-CAAACTGGGATTAGATACC-3′) and 12S-9R (5′-AGAACAGGCTCCTCTAG-3′). The relatively short target sequence was selected to increase detection of degraded DNA in partially digested blood meals [54]. All DNA samples and negative controls from DNA extractions were included in the blood meal analysis. Sequencing libraries were prepared using a two-step PCR protocol in which the target region was first amplified, then amplicons converted into Illumina-compatible libraries through the addition of indexed adapter sequences. Resulting libraries were sequenced using the Illumina MiSeq platform using a PE300 kit, generating 2×300bp paired-end reads. Raw sequence data were processed by the University of Delaware Center for Bioinformatics and Computational Biology. Sequence analysis was conducted using VSEARCH v2.30.1 [55]. Paired-end reads were merged, trimmed, and filtered to remove low-quality or off-target sequences. Amplicon sequence variants (ASVs) were then generated from the remaining sequences.

To address index hopping or other sequencing artifacts, we implemented a sample percentage threshold following Drake et al. [56] consisting of removing any ASV that represented less than 2% of the total read count for that sample. To address any possible DNA contamination, negative controls from each batch of DNA extractions were sequenced with specimen DNA. Any ASVs that amplified in the negative controls were removed from samples corresponding to those controls. After filtering out ASVs from negative controls and those that represented less than 2% of each sample’s total reads, ASVs were manually reviewed and given a final taxonomic assignment by querying each sequence against the National Center for Biotechnology Information (NCBI) nucleotide database using the Basic Local Alignment Search Tool (BLAST; [57,58]). We selected the highest-scoring BLAST match, with a minimum sequence identity of 95%, that was regionally (i.e., found in the Mid-Atlantic, US) and biologically (i.e., a terrestrial vertebrate) plausible. If all matches were implausible or under 95% sequence identity, the ASV was excluded. Sequences that corresponded to nuclear mitochondrial DNA segments (NUMTs) as well as arthropod ASVs that remained after quality filtration were also excluded.

### Human blood meal analysis

Given the epidemiological importance of human blood meals in a region that is not considered Chagas endemic, we subjected each specimen to two other human-specific assays in addition to the 12S deep sequencing: (i) an immunochromatographic rapid test targeting a human blood protein and (ii) a second deep sequencing assay targeting a human-specific gene sequence. For the rapid test, we used Rapid Stain Identification of Human Blood (RSID) kits (Independent Forensics). The RSID test can detect as little as 1μL of human blood by targeting the human red blood cell transmembrane protein glycophorin A. The test was originally developed for use in forensics and is human-blood-specific [59] but validated for use with triatomine gut contents [60]. To carry out the RSID test, we used a sterile scalpel to make a small incision in the ventral abdomen of each *T. sanguisuga* specimen and inserted a sterile cotton-tipped applicator through the opening to collect gut material. We then immersed the tip of each applicator into 200μL of RSID Universal Buffer and continued the protocol according to the manufacturer’s instructions.

Our second human-specific assay was a next generation sequencing procedure targeting a 228bp fragment of the human mitochondrial cytochrome c oxidase subunit I (COI) gene using the cox I primers (forward: 5′-TTCGGCGCATGAGCTGGAGTCC-3′, reverse: 5′-TATGCGGGGAAACGCCATATCG-3′; [61]). The assay was performed at the University of Delaware Sequencing and Genotyping Center where Illumina sequencing was carried out as described above for the 12S assay. Raw sequence data were processed using QIIME2, with DADA2 used for quality filtering [62,63]. Following bioinformatic processing, we filtered COI sequence read data to mitigate possible DNA contamination and sequencing artifacts. As this assay was species-specific, we anticipated having blank samples and determined a total percentage threshold was more appropriate than a sample percentage threshold for these data [56]. For each sample, we removed occurrences of any ASV that represented less than 0.002% of the total read count, a threshold of 485bp which in practice eliminated all read counts less than or equal to 15bp, which was within the range recommended by Drake et al. [56]. To mitigate effects of DNA contamination, we implemented a maximum contamination filter, removing any occurrence of an ASV that had amplified in a corresponding negative control, if read counts were less than those seen in the negative control. Following this process, we manually reviewed each ASV and determined final taxonomic assignments by querying each ASV sequence against the NCBI nucleotide database using BLAST. We retained only ASVs with a full alignment length of 184bp and a 100% identity match to a human mitochondrial sequence.

### Data analysis

All analyses were carried out in the R statistical computing environment [64]. We tested for associations between *T. cruzi* infection status, *T. sanguisuga* life stage (adult or nymph), and study site LUI using Fisher’s exact tests and a significance threshold of 0.05.

We tested for differences in blood meal host species between adult and nymphal stages using a permutational multivariate analysis of variance (PERMANOVA) run with the R package *vegan* [65,66]. Sequence read data were collapsed into binary presence-absence variables indicating detection of each host species within individual specimens, excluding specimens with no identifiable vertebrate blood meals. These data were used to calculate a Jaccard dissimilarity matrix on which we conducted the PERMANOVA. Statistical significance was assessed using 999 permutations. To identify individual blood meal host species associated with adult and nymphal stages, we performed an indicator species analysis following Dufrêne and Legendre (1997). We conducted this analysis using the group-equalized indicator value (IndVal.g) index with 999 permutations [67]. To deal with uneven collection of adults and nymphs between sampling sites (several sites yielded either adults or nymphs but not both), we included study site as a covariate in the PERMANOVA and also repeated the analysis with specimens only from sites where both nymphs and adults were collected.

### Blood meal host network analysis

To evaluate potential *T. cruzi* transmission pathways within the blood meal host community, we constructed a weighted host co-occurrence network from bloodmeal data. Sequence read data were collapsed into binary presence-absence variables indicating detection of each host species within individual specimens, excluding specimens with no identifiable vertebrate blood meals. For specimens containing multiple host species, we converted blood meals into pairwise combinations of co-occurring hosts. Each node in the network represented a different vertebrate host species and edges (the connections between nodes) represented co-occurrence of those species within the same *T. sanguisuga* specimen. Edge weights reflected the frequency of co-occurrence across all specimens. Network structure was evaluated using three node-level centrality metrics calculated with the R package *igraph* [68] with degree (*k*) as the sum of the number of nodes to which a given node was directly connected (i.e., the number of other species co-occurring with a given species in the host network). Strength (*s*) was the cumulative frequency of direct connections between a given node and other nodes (i.e., the sum of total co-occurrences of a given species with all other species in the host network). Eigenvector centrality (EC) measured the relative influence of a node within the network, taking into account the degree of the node as well as the degree of the nodes to which it was connected [69]. Degree and strength were strongly influenced by sample size, but eigenvector centrality was standardized to a maximum value of 1, allowing for the comparison of nodes between different networks. A separate network using only *T. cruzi*-infected specimens was constructed to evaluate whether patterns of host connectivity by *T. cruzi*-infected bugs differed from the overall network, and to identify potentially important *T. cruzi* host species.

## Results

### *Triatoma sanguisuga* sampling

We collected a total of 151 *Triatoma sanguisuga* specimens from 25 locations in Delaware and five locations in eastern Maryland for a total of 30 locations with *T. sanguisuga* (Fig 1). Active sampling (light sheets and manual searching) yielded 131 of the 151 specimens, which came from 21 of the 28 sample sites. The remaining 20 specimens were community submissions from 10 locations. The bucket traps did not yield any specimens. All three Delaware counties (New Castle, Kent, Sussex) as well as Cecil and Harford counties in Maryland (community submissions) were represented in our collection. All community submissions were found between the months of June and September. Seven specimens were found inside human homes (Table 2). All community submissions that were found in or around human infrastructure were adult specimens.

**Figure 1.**
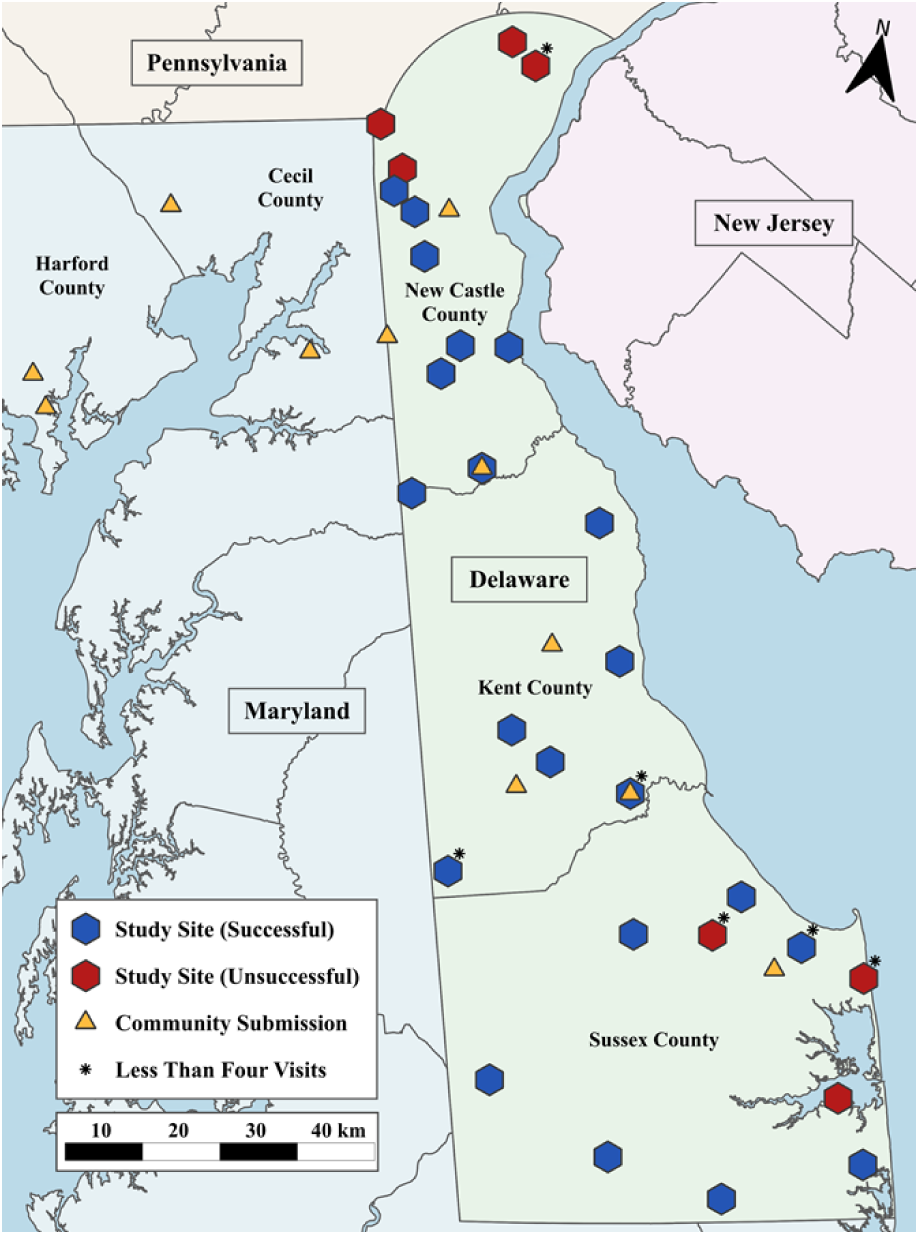
*Triatoma sanguisuga* collection locations. Successful sites indicate that at least one *T. sanguisuga* specimen was collected; unsuccessful sites were sampled but no specimens were collected. All study sites were sampled a minimum of four times except those with an asterisk.

**Table 2.** Locations in and around human homes and businesses where *T. sanguisiga* was found. All samples were in Delaware and eastern Maryland.

|  | **Location** | **Month** | ***T. cruzi*** | **Human blood** |
|---|---|---|---|---|
| **Indoors** | Bathroom ceiling | June | - | - |
|  | Windowsill (room unspecified) | June | + | - |
|  | Kitchen sink | July | + | - |
|  | Laundry room | July | - | - |
|  | Commercial office | August | + | - |
|  | Living room fireplace | August | + | + |
|  | Bathtub | September | - | - |
|  | Living room curtains | September | - | - |
| **Outdoors** | Back patio of private home | July | + | + |
|  | Backyard of private home | July | + | - |
|  | Gas station (artificial light) | July | + | - |
|  | Gas station (artificial light) | July | - | - |
|  | Gas station (artificial light) | July | + | - |
|  | Front entrance of building | July | - | - |
|  | Loading dock behind building | July | + | - |
|  | Pool skimmer | July | + | - |

*Triatoma sanguisuga* life stage consisted of 116 (76.8%) nymphs and 35 (23.2%) adults (19 males and 16 females). Specimens were found at sites of all three land use intensities (LUI), with the highest number (82/151; 54.3%) found at medium LUI sites and the lowest number (40/151; 27%) found at low LUI sites. The majority of specimens collected at high LUI sites (19/29, 65.5%) were adults, while most specimens collected at medium (70/82, 85.4%) and low (36/40, 90.0%) LUI sites were nymphs. As mentioned, snags were the most reliable source of *T. sanguisuga* specimens, accounting for 78.1% (118/151) of specimens collected including all of the nymphs (N=116) and 6/35 (17.1%) of the adults. These specimens came from 49 snags across 16 study sites. We collected up to 11 specimens from a single snag, while 24 other snags (49.0%) yielded only a single specimen.

### *Trypanosoma cruzi* infection

Trypanosomes were observed in the excrement of 9 of 129 (7%) *T. sanguisuga* specimens examined via direct microscopic observation (Table S2). A total of 23 specimens, including the 9 individuals in whose excrement trypanosomes were observed, were positive for *T. cruzi* via RT-PCR. All positive specimens harbored *T. cruzi* DTU TcIV only. *Trypanosoma cruzi* infection status differed significantly between life stages; 19 of 35 adult specimens (54.3%) were infected with *T. cruzi* compared to 4 of 115 nymphs (3.5%). Odds of infection in adult *T. sanguisuga* were 31.7 times higher than in nymphs (Fisher’s exact test; OR = 31.7; p < 0.001; Fig. 2), suggesting that regional *T. cruzi* infection prevalence in *T*.

**Figure 2.**
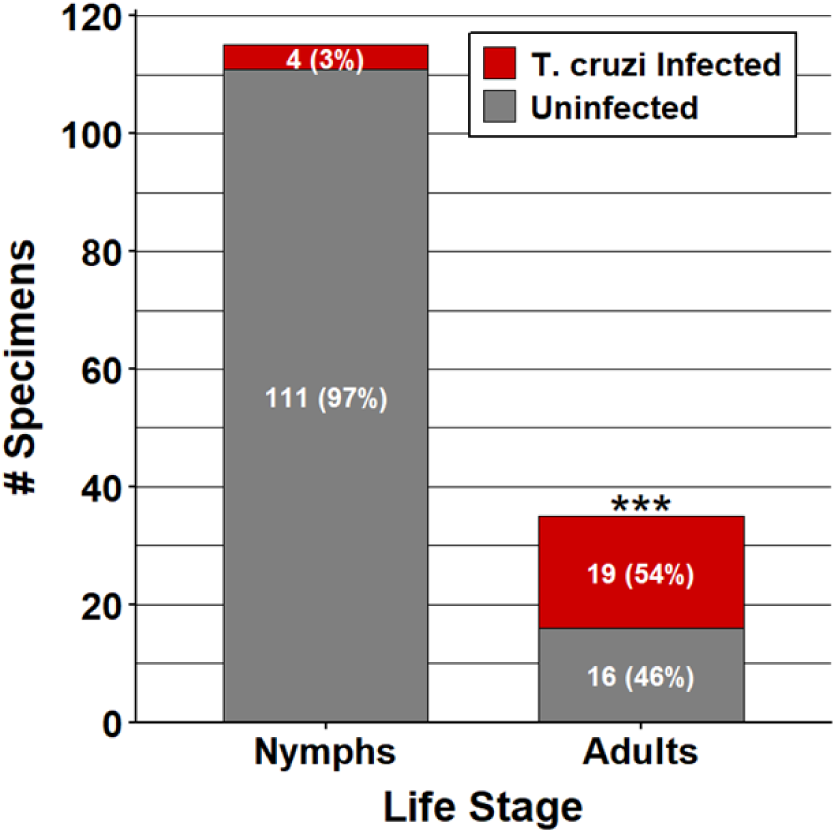
*T. cruzi* infection in *T. sanguisuga* by life stage. ***p<0.001. *sanguisuga* may be stage structured.

*Trypanosoma cruzi* infection status differed significantly between study site LUI (Fisher’s exact test; p < 0.001) with highest prevalence found in high LUI sites (13/29, 44.8%), which was probably an effect of the number of adult specimens (19/29) found at high LUI sites. Among the 20 community submissions, 10 (50%) were infected with *T. cruzi*.

### Blood meal analysis

Sequence data for the 12S assay were successfully obtained from 147 of 151 specimens (97.4%) and we identified 24 vertebrate host species from 95 *T. sanguisuga* specimens comprising 13 mammal species, three reptile species, four amphibian species, and four avian species (Fig. 3, Table S2.1). The highest number of total reads and blood meal occurrences were from two species, eastern gray squirrel (*Sciurus carolinensis*; detected in 32 specimens and accounting for 26.3% of all reads) and five-lined skink (*Plestiodon fasciatus*; detected in 29 specimens and accounting for 28.1% of all reads). We identified an average of 1.5 host species (SD = 0.87) per *T. sanguisuga* specimen, with multiple blood meal hosts detected in 33 specimens, including three or more species in 10 specimens. The maximum number of host species identified in a specimen was five. A list of NCBI BLAST alignment results for each ASV is in Table S2.2.

**Figure 3.**
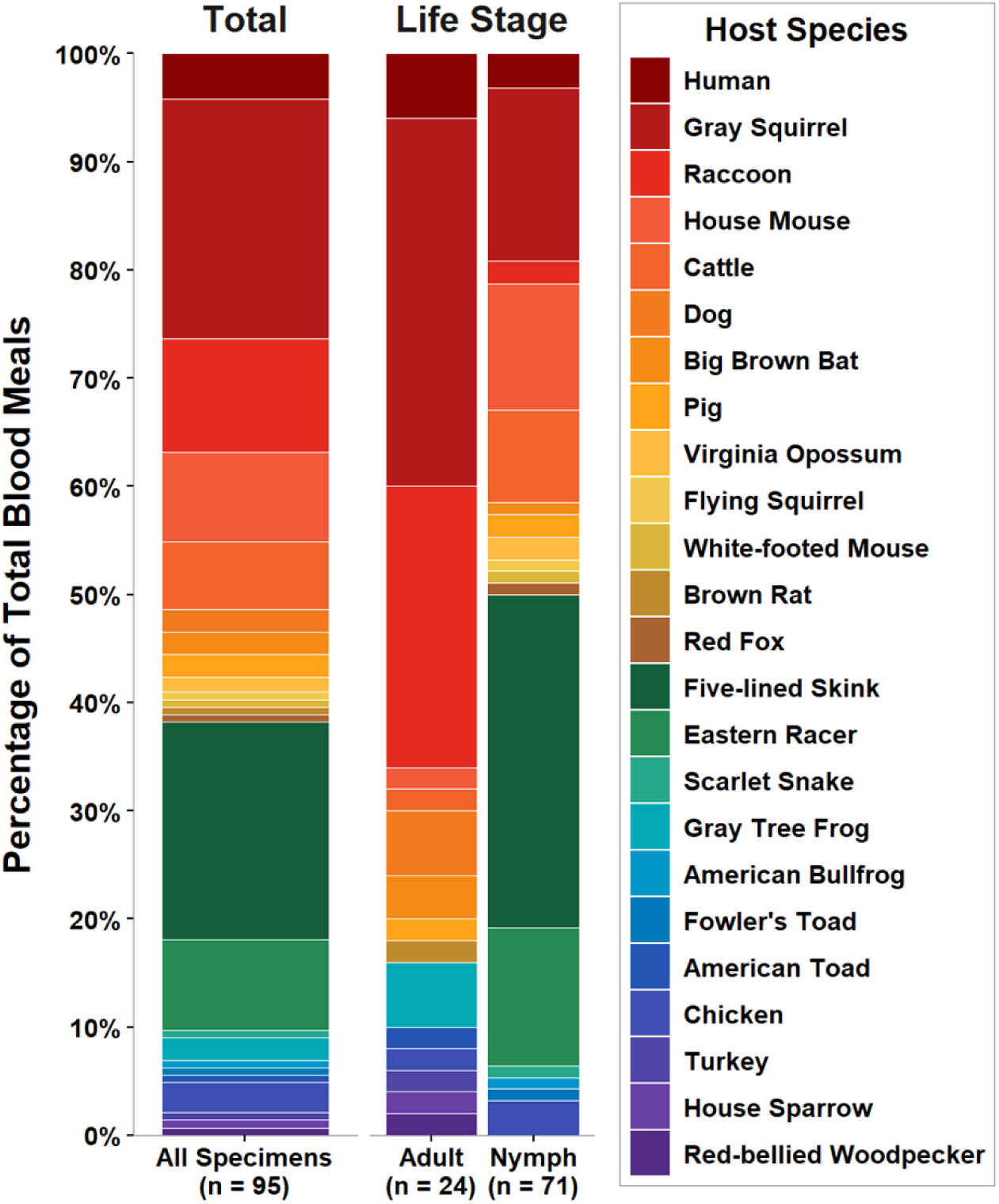
Blood meal host species identified in *T. sanguisuga in* Delaware and eastern Maryland. *Trypanosoma cruzi* competent hosts (i.e., mammals) are shown in warm tones (reds, oranges, yellows). Refractory hosts (i.e., reptiles, birds, and amphibians) are shown in cool tones (greens, blues, and purples).

### Human blood meals

As mentioned, we detected human blood meals using three different methods: RSID rapid tests targeting a human blood cell surface protein and two Illumina sequencing assays targeting a 145bp universally conserved vertebrate 12S sequence and a 228 bp human-specific COI sequence. RSID tests detected human blood in 19/133 (14.3%) *T. sanguisuga* specimens tested with the 12S sequencing assay amplified human DNA in 38/151 (25.2%) *T. sanguisuga* specimens, represented by one ASV after data filtration (Table S2.3). In the human-specific COI sequencing assay, human DNA was detected in 50/151 (33.1%, Table 3) *T. sanguisuga* specimens, represented by five ASVs after data filtration (Table S2.3).

**Table 3:** Human blood meal results by detection method. NA= RSID test not conducted.

|  |  | **RSID** |  |  |  |
|---|---|---|---|---|---|
|  |  | **Positive** | **Negative** | **NA** | **Total** |
| **Sequencing Assays** | **12S + / COI +** | 1 (0.7%) | 15 (9.9%) | 2 (1.3%) | 18 (11.9%) |
|  | **12S + / COI –** | 2 (1.3%) | 15 (9.9%) | 3 (2.0%) | 20 (13.2%) |
|  | **12S – / COI +** | 5 (3.3%) | 25 (16.6%) | 2 (1.3%) | 32 (21.2%) |
|  | **12S – / COI –** | 11 (7.3%) | 59 (39.1%) | 11 (7.3%) | 81 (53.6%) |
| **Total** |  | 19 (12.6%) | 114 (75.5%) | 18 (11.9%) | 151 (100%) |

### Negative controls

In the 12S assay, human DNA amplified in two negative controls from the extraction step, which corresponded to 19 samples; all human reads were excluded from those 19 samples. The other 10 negative extraction controls did not amplify human DNA (corresponding to 132 samples). Human DNA did not amplify in any extraction control in the COI assay, but one human-identified ASV amplified at a relatively low level in one of two no-template controls introduced during Illumina library preparation. This ASV was excluded from samples with read counts lower than the no-template control (4756bp; 70 samples excluded; 32 samples preserved). In the RSID reaction, all negative control cassettes were negative. The RSID test was human blood specific, meaning it did not react to other human bodily fluids such as aerosolized saliva or mucous, making it less likely to be contaminated than DNA-based methods.

The three detection methods were discordant; a single specimen (1/151, 0.7%) tested positive for human blood by all three methods; this specimen was a community submission that had been found on the leg of a table in the backyard of a private home. In contrast, 81/151 (53.6%) specimens were positive for human blood by at least one of the three detection methods. Due to these discordant results, we took a conservative approach for inclusion of human blood meals when analyzing determinants of blood meal host species and host networks (below). For these analyses, we included human blood meals only from specimens (N = 6) that were positive for human blood on both human specific assays (COI and the RSID tests).

### Determinants of blood meal host species

When investigating determinants of host species identified in *T. sanguisuga* blood meals, we found that geographic location and insect life stage explained 35.9% and 1.9% of the variation in the data, respectively (PERMANOVA; study site: R² = 0.359, p < 0.01; life stage: R² = 0.019, p < 0.05). Four host species were significantly associated with adult *T. sanguisuga*: gray squirrel (IndVal.g = 0.74, p < 0.01), raccoon (IndVal.g = 0.72, p < 0.01), dog (IndVal.g = 0.35, p < 0.05), and gray tree frog (IndVal.g = 0.35, p < 0.05). Five-lined skink was a significant indicator of nymphs (IndVal.g = 0.64, p < 0.01).

When visualizing these data in a principle coordinates analysis plot using the Jaccard dissimilarity index for life stage, the first two principal coordinates (PC1 and PC2) collectively described 50.2% of the variation in the data (Fig. 4). The host species fed on by adults and nymphs did overlap, but the centroids of these groups were distinct, due largely to some nymphs exhibiting host-use profiles (i.e. feeding on reptiles) that were not present in adult specimens. These results suggest that there is not a complete partition between adult and nymph host communities but rather that adults restrict their host-use to a subset of the species fed on by nymphs.

**Figure 4.**
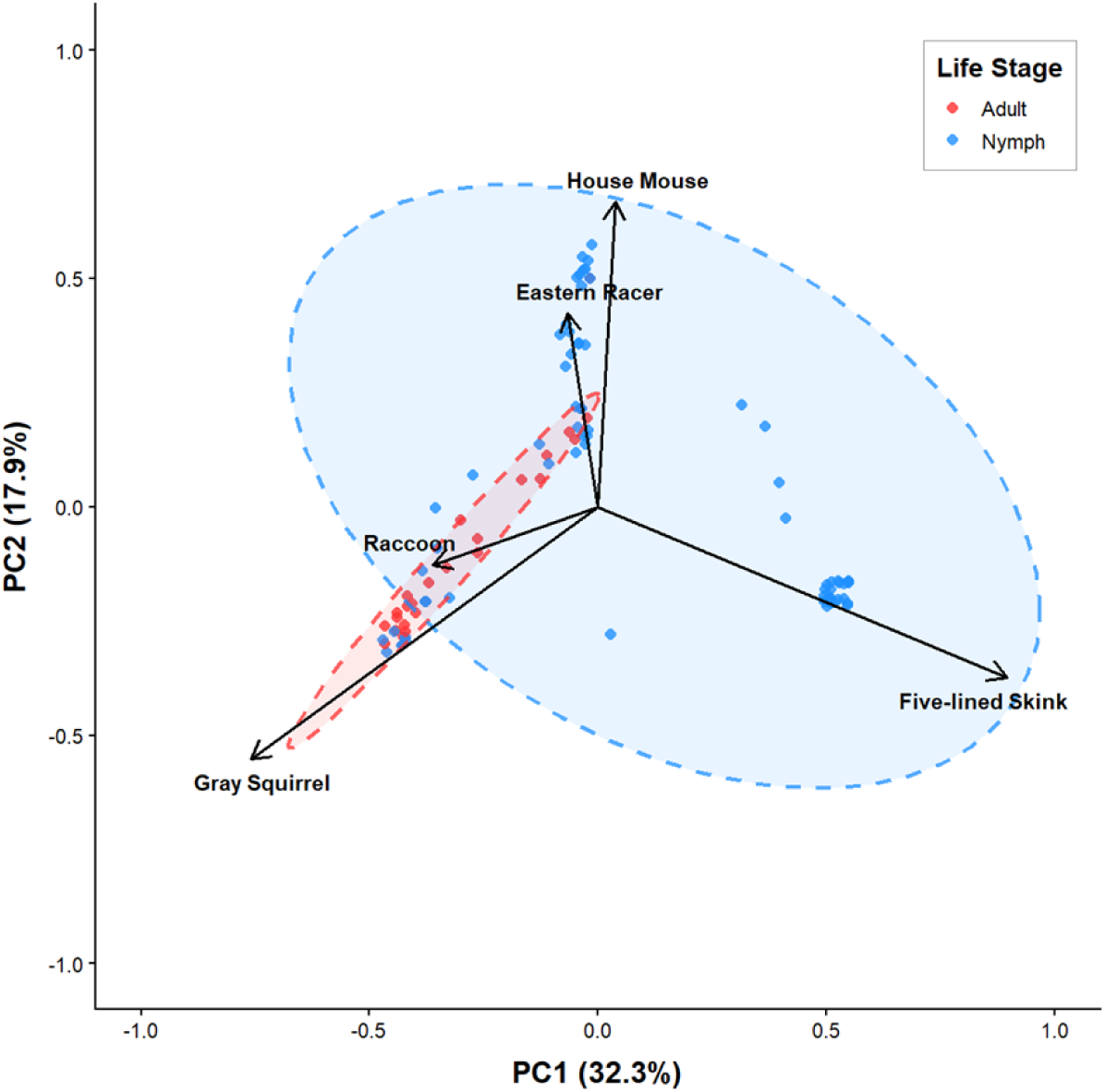
PCoA plot of blood meal host species fed on by *Triatoma sanguisuga* using the Jaccard dissimilarity index for life stage.

To check if the apparent stage-structured host species preferences findings were confounded by inherent variation in host species diversity and abundance between study sites, we repeated the PERMANOVA analysis with a subset of our data comprising only the specimens collected at sites in which both life stages were collected (5 sites; 13 adults and 27 nymphs). We again found geographic location and insect life stage to be significant, with geographic location reduced to explaining 18.0% of the observed variation and life stage increasing to 4.8% of the observed variation (PERMANOVA; study site: R^2^ = 0.180, p < 0.05; life stage: R^2^ = 0.048, p < 0.05). Raccoon was the only significant indicator of adult specimens (IndVal.g = 0.74, p < 0.01), and no host species were associated with nymphs. These results suggest that while host associations vary significantly between study sites, the difference between adult and nymph blood meal host feeding patterns is a significant effect independent of local host availability.

### Host networks

The relative importance of species within the vertebrate host network was variable as measured by three node-level centrality metrics (Table S2.4). As mentioned, degree (*k*) quantified the number of other species connected to a given species by blood meal co-occurrence, strength (*s*) quantified the cumulative frequency of those connections, and eigenvector centrality (EC) quantified the relative influence of a species within the network. Gray squirrel was the node with the highest score in all centrality metrics (*k* = 14, *s* = 31, EC = 1.00). Racoon was the second-highest scoring species (*k* = 9, *s* = 23, EC = 0.92) and humans were third highest (*k* = 7, *s* = 13, EC = 0.55; Fig. 5, Table S2.4). The network structure of *T. cruzi*-infected specimens was similar, with gray squirrel scoring highest (*k* = 9, *s* = 19, EC = 1.00) followed by raccoon (*k* = 6, *s* = 16, EC = 0.98). Some host species were absent from the *T. cruzi* infected host network, including all reptiles. Other host species were more central (gray tree frog, ΔEC = +0.15) or more peripheral (big brown bat, ΔEC = -0.15) in the infected network.

**Figure 5.**
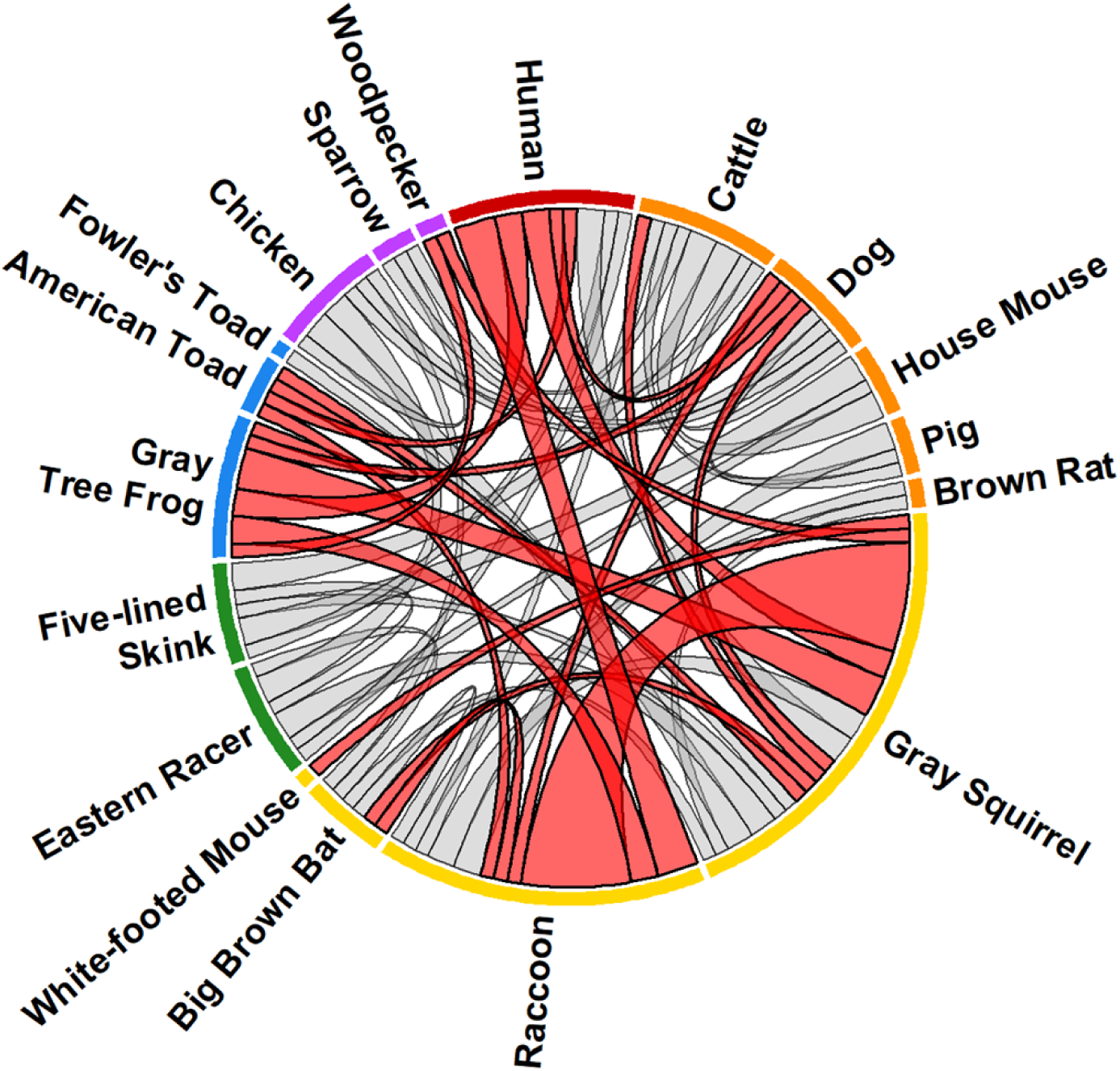
Host co-occurrence network. Nodes represent host species; node size is proportional to number of detections. Node color: red = human, orange = domestic mammal, yellow = sylvatic mammal, green = reptile, blue = amphibian, purple = bird. Linked nodes indicate co-occurrence of species within *T. sanguisuga* specimens. Link width is proportional to frequency of co-occurrence. Red links indicate *T. cruzi*-infected specimens. Specimens for which just one blood meal host species was detected are not included in the network.

## Discussion

Here we carried out the first investigation of *Triatoma sanguisuga,* vector of Chagas disease agent *T. cruzi*, in Delaware and two counties in eastern Maryland. We found that *T. sanguisuga* is widely distributed throughout the region across different land use intensities, although nymphs were found only in standing dead trees (‘snags’). We discovered that *T. cruzi* prevalence in *T. sanguisuga* is stage structured, with adults being over 30 times more likely to be infected with *T. cruzi* than nymphs, which is likely a reflection of differences in host use between adults and nymphs, as evidenced in our blood meal analysis. Human exposure to triatomine bugs probably occurs, as 15 specimens were found in or around human homes and businesses, although discordant blood meal assay results made the level of exposure impossible to quantify. Nevertheless, this work has provided valuable, region-specific ecological information on an understudied local vector species and serves as a first step toward making evidence-based assessments of the local *T. cruzi* transmission risk for both humans and their animals.

### *T. sanguisuga* is distributed across a variety of Delaware landscapes south of I-95

Although *Triatoma sanguisuga* is native to the eastern seaboard states, the species was not reported in Delaware by the CDC until 2019 [70] and its geographic distribution in the state was unknown. We found *T. sanguisuga* in residential, recreational, and forested landscapes in all three Delaware counties as well as two neighboring counties in Maryland. We identified Interstate 95 (I-95) as a possible northern limit to the species’ distribution in Delaware based on repeated unsuccessful sampling of three sites situated just north of this boundary. These sites align with geological (fall line) and topographical (coastal plain to piedmont) transitions zones that are accompanied by subtle changes in vegetation and climate that could play a role in restricting *T. sanguisuga*’s distribution north of I-95. However, this boundary is also the beginning of the densely urbanized Wilmington metropolitan area in Delaware, and effects of urbanization could be a limiting factor. The coincidence of changes in urbanization and topography make it difficult to identify the underlying mechanism of the northern limit of the species in Delaware. In addition, we did not sample a narrow slice of coastal plains that runs north of I-95, following the northeastern coast of Delaware and continuing up along the Delaware River through Philadelphia, PA and up to Trenton, NJ. This small corridor could still serve as *T. sanguisuga* habitat and studies of the region are needed. Notably, an adult female *T. sanguisuga* specimen collected in Philadelphia in 1931 is housed in the Smithsonian Entomological collection (Catalog# USNMENT01240718), confirming that the species was once there. Finally, records of both *T. sanguisuga* and the northwesternmost triatomine species (*Hospesneotomae protracta*) stop just below the 42^nd^ parallel N across the entire north American continent, suggesting that broader scale drivers of triatomine distribution, such as climate patterns, may be at play as well [71,72].

### Snags are an important local habitat for *T. sanguisuga*

In the southwestern US, *T. sanguisuga* is associated with packrats (*Neotoma spp*), which cohabitate with triatomine bugs in packrat middens [73]. Reports of habitat used by *T. sanguisuga* in regions without *Neotoma* species are sparse but consistent, dating to 1869 and spanning several US states, although no reports exist from the northeastern or Mid-Atlantic, US. These reports describe *T. sanguisuga* under loose tree bark in the subcortical space of dead trees and hollow stumps [74–80]. Ryckman suggested that *T. sanguisuga* in the eastern US inhabited large hollow trees [81,82], which is what we found - standing dead trees (aka ‘snags’) were a reliable collection location for *T. sanguisuga* in DE; every nymph collected in this study (N=116) was collected from a snag, as well as six adults. Our findings suggest that snags are an important local habitat for *T. sanguisuga* in Delaware.

Snag cavities provide an insulated environment that has been found to maintain higher autumn and winter temperatures than the surrounding environment [83], and they are likely used by *T. sanguisuga* for overwintering, which has been reported previously in other regions [74,77,78,80]. We found nymphs under the bark of snags throughout the summer as well, suggesting that snags might also serve as *T. sanguisuga* breeding, oviposition, and juvenile habitat. While many triatomine species oviposit continuously, egg laying in *T. sanguisuga* is cyclical, occurring in the wild only between May and September [36,84]. Future studies should clarify if snags are the sole habitat for *T. sanguisuga* in the greater Mid-Atlantic as well, or merely one of several local refugia that the species inhabits.

### *T. sanguisuga* in Delaware appears to be a generalist feeder

We found that *T. sanguisuga* in Delaware is a generalist that feeds on an array of vertebrate species across different land use types, which aligns with blood meal analysis studies of *T. sanguisuga* in the southern US [13,16,25,85,86]. Common blood meal host species for *T. sanguisuga* in this study and in previous works were gray squirrel (*Sciurus carolinensis*), raccoon (*Procyon lotor*), and skinks (*Plestiodon spp*). In addition, we identified ten new blood meal host species that were not reported for *T. sanguisuga*, comprising five mammal species (brown rat, big brown bat, southern flying squirrel, red fox, white-footed mouse), two amphibian species (American bullfrog, gray treefrog), one reptile (scarlet snake), and two birds (house sparrow, red-bellied woodpecker), expanding its generalist feeding repertoire (Fig. 5, Table S2.2).

### *Stage-structured* feeding, *T. cruzi* prevalence, and habitat partitioning

We found that adult *T. sanguisuga* were 30 times more likely to be infected with *T. cruzi* than *T. sanguisuga* nymphs. This stage-structured *T. cruzi* prevalence may be due to the stage-structured feeding pattern we observed wherein *T. sanguisuga* adults specialized on mammals, which are *T.* cruzi competent, while nymphs fed on a wider range of taxa of which over 50% were refractory hosts of *T. cruzi* (i.e., not mammals). More specifically, *T. sanguisuga* nymphs fed heavily on the five-lined skink (which nests in the same habitat used by the nymphs), while adults did not feed on any reptiles. The authors of the only other study of *T. sanguisuga* nymphal blood meals also observed a difference in host use between nymphs and adults [13].

The stage structured patterns we observed in *T. sanguisuga* in Delaware might also extend to its local *T. sanguisuga* habitat use. Many of the animals on which *T. sanguisuga* nymphs fed nest in subcortical spaces under the peeling bark of snags. Adults on the other hand, fed more on canopy dwelling species, suggesting a possible stage-structured vertical partitioning of arboreal habitats: nymphs, which are wingless, inhabit tree trunks, feeding opportunistically on locally available hosts; adults, which have functional wings, move upwards to feed on mammals that nest in the canopy. This vertical partitioning of tree habitats was described for *T. sanguisuga* by Raymond Ryckman in his seminal works on the Triatominae, [73,81], although he described the behavior for geographically separate populations of *T. sanguisuga* (Florida vs. other states east of the Mississippi river). Since we did not directly sample the canopy but rather inferred canopy use by adult *T. sanguisuga* through blood meal host species, future studies should directly sample the living canopy to clarify whether or not the stage-structured host use is a result of vertical partitioning of tree habitats between *T. sanguisuga* life stages.

### Raccoons as possible *T. cruzi* reservoirs in Delaware

The coupling of raccoon blood meals and high *T. cruzi* prevalence in adult *T. sanguisuga* suggest that raccoons may serve as a *T. cruzi* reservoir species in Delaware. Raccoons are relatively well-studied *T. cruzi* hosts, and they have been implicated as *T. cruzi* reservoirs in several other states [87]; in the Mid-Atlantic, raccoons have been found infected with *T. cruzi* in Virginia [33–35] and Maryland [88,89]. Additionally, raccoons are thought to be associated with *T. cruzi* DTU TcIV [87,90], which was the only *T. cruzi* type detected in our infected *T. sanguisuga* specimens. Studies of *T. cruzi*-infected raccoons found that they sustained prolonged parasitemia [91] and were highly infectious to vectors. A 2017 review of *T. cruzi* in US wildlife found that raccoons had the highest aggregate infectiousness index and overall *T. cruzi* infection prevalence (36.4%) of 26 wildlife species analyzed [87]. Our data suggest that the raccoon’s role as an important *T. cruzi* reservoir extends to Delaware, but *Trypanosoma cruzi* infection in vertebrate hosts in Delaware should be measured to clarify the role of raccoons as well as other *T. cruzi* host species.

### Human blood feeding

Although we employed multiple precautionary measures at every step of our three human blood meal detection assays, test results were discordant. As a result, we cannot confidently estimate the frequency of human blood feeding by *T. sanguisuga* in Delaware. We suspect that the discordance is due to biological differences between assay targets, comprising sialoglycoproteins in the RSID tests and mitochondrial DNA in the deep sequencing assays.

These molecules differ greatly in their biochemical and structural properties as well as in their distribution and abundance in human tissues, which could contribute to differences in sensitivity and specificity between assays. To our knowledge, this is the first study to use RSID and DNA-based human blood meal detection in tandem. Previous studies compared multiple DNA-based assay targets to detect human blood meals, and these studies also produced discordant results [92–94]. Stevens et al. (2012) conducted a blood meal analysis on triatomines using both the cytB and 12S primers, and found that the results of each assay identified different numbers of host species, different numbers of blood meals, and different haplotypes [95]. This outcome is generally considered positive because combining multiple loci broadens the scope of the assay and improves host species detection [96]. However, in the case of triatomine bugs in a region considered non-endemic for Chagas disease with an unknown risk for *T. cruzi* exposure, our objective was not to maximize the detection of human blood meals, but rather to minimize false positives and gain an accurate view of *T. sanguisuga* contact with humans. Our results indicate that controlled studies are needed to determine relative sensitivity thresholds between RSID and DNA-based human blood meal detection methods and a gold standard should be determined. Our findings also highlight the inherent uncertainty in any type of human blood detection assay, even when all possible precautions are taken to conserve the integrity of a sample, illustrating that these data must always be interpreted and communicated stringently.

### Epidemiological relevance of *T. sanguisuga* in Delaware

Studies of human blood feeding by *T. sanguisuga* in in Texas, Louisiana, and Florida found that between 22% to 76% of specimens had fed on humans [13,25,97]. While we cannot confidently estimate the biting rate of *T. sanguisuga* in Delaware, we can say that seven of our 151 (4.6%) specimens were collected inside human homes, including bedrooms. In addition, one adult specimen was collected inside an office, and seven others were collected in yards or exterior walls of homes and businesses. All *T. sanguisuga* encounters occurred during the summer months. Twelve (80%) of these 15 specimens tested positive for human blood on at least one assay and nine (60%) were infected with *T. cruzi*. Collectively these results indicate that *T. cruzi-*infected *T. sanguisuga* do occasionally invades homes and other buildings in Delaware where humans are present.

The occasional intrusion of *T. sanguisuga* in human homes and yards is not unexpected; sporadic reports exist dating back to 1869 when Walsh and Riley reported *T. sanguisuga* in southern Illinois biting humans in their beds [74]. However, reports of *T. sanguisuga* nymphs inside homes, the hallmark of synanthropic triatomine species, have not occurred in Delaware and are rare in other part of the US as well [98]. Together, these findings suggest that public health efforts should be directed toward education on preventing *T. sanguisuga* intrusion into homes and other residential or commercial structures as well as how to safely handle and dispose of triatomine bugs. Additionally, there is a lack of awareness in the medical community about the existence of a native *T. cruzi* vector species in Delaware, which has resulted in local citizens being dismissed by their physicians when seeking advice or testing for *T. cruzi* [17]. Thus, a second important public health step would be to educate both the medical community and the community at large on steps that can be taken in response to a suspected triatomine bug bite.

### Conclusion

Here we presented findings from the first ecological investigation of *T. sanguisuga* and *T. cruzi* in the Mid-Atlantic US, revealing key aspects of *Triatoma sanguisuga* in temperate environments, including its habitat, phenology, bloodmeal hosts, and *T. cruzi* infection. Our results suggest that *T. sanguisuga* in Delaware and eastern Maryland breeds in forested areas where it predominantly feeds on wildlife but forages in residential and commercial areas in the summer months. Observations of adult bugs in and around human infrastructure highlight the need for public education and awareness about *T. sanguisuga* to increase public health preparedness in the Mid-Atlantic, US.

## Acknowledgements

The authors would like to thank the following organizations for allowing the collection of Triatomine bugs on their properties: U.S. Fish and Wildlife Service Coastal Delaware National Wildlife Refuge Complex, Delaware Department of Natural Resources and Environmental Control (DNREC), Delaware Division of Fish and Wildlife, Delaware Wild Lands, Delaware Forest Service, Delaware National Estuarine Research Reserve (DNERR), and the city of Odessa, DE. The authors are also grateful to the community members and students who helped to with the project including Anthony Alvarado, Joshua Bast, Brian Bahder, Kevin Harkins, Solomon Hendrix, Nathan Levia, Richard Rice, and Sarah Sharp. This work received funding from DE-INBRE, which is supported by NIH NIGMS IDeA Program Grant P20GM103446 and the State of Delaware as well as the USDA National Institute of Food and Agriculture, Hatch Multistate project NE2443.

